# Open-Source Food: Nutrition, Toxicology, and Availability of Wild Edible Greens in the East Bay

**DOI:** 10.1101/385864

**Authors:** Philip B. Stark, Daphne Miller, Thomas J. Carlson, Kristen Rasmussen de Vasquez

**Affiliations:** Department of Statistics, University of California, Berkeley, California, United States of America; School of Public Health, University of California, Berkeley, California, United States of America; Department of Family and Community Medicine, University of California, San Francisco, California, United States of America; Department of Integrative Biology, University of California, Berkeley, California, United States of America; Department of Nutrition and Toxicology, University of California, Berkeley, California, United States of America

## Abstract

**Significance:** Foraged leafy greens are consumed around the globe, including in urban areas, and may play a larger role when food is scarce or expensive. It is thus important to assess the safety and nutritional value of wild greens foraged in urban environments.

**Methods:** Field observations, soil tests, and nutritional and toxicology tests on plant tissue were conducted for three sites, each roughly 9 square blocks, in disadvantaged neighborhoods in the East San Francisco Bay Area in 2014–2015. The sites included mixed-use areas and areas with high vehicle traffic.

**Results:** Edible wild greens were abundant, even during record droughts. Soil at some survey sites had elevated concentrations of lead and cadmium, but tissue tests suggest that rinsed greens of the tested species are safe to eat. Daily consumption of standard servings comprise less than the EPA reference doses of lead, cadmium, and other heavy metals. Pesticides, glyphosate, and PCBs were below detection limits.

The nutrient density of 6 abundant species compared favorably to that of the most nutritious domesticated leafy greens.

**Conclusions:** Wild edible greens harvested in industrial, mixed-use, and high-traffic urban areas in the San Francisco East Bay area are abundant and highly nutritious. Even grown in soils with elevated levels of heavy metals, tested species were safe to eat after rinsing in tap water. This does not mean that all edible greens growing in contaminated soil are safe to eat—tests on more species, in more locations, and over a broader range of soil chemistry are needed to determine what is generally safe and what is not. But it does suggest that wild greens could contribute to nutrition, food security, and sustainability in urban ecosystems. Current laws, regulations, and public-health guidance that forbid or discourage foraging on public lands, including urban areas, should be revisited.

## Introduction

Diets that include a wide array of plant-based foods are associated with lower rates of chronic disease and better health outcomes [1–3]. Edible wild and feral plants—edible weeds—are an abundant but generally overlooked food source with the potential to contribute to dietary diversity and vegetable intake, especially in areas with limited access to fresh produce.

Edible weeds, which require neither cultivation nor intentional watering, are generally abundant in farms, gardens, parks, yards, sidewalks, and medians, on both private and public land: anywhere there has not been a concerted effort to eradicate them. Some are native to their habitat, but many are nonnative feral species that were once cultivated deliberately but have since colonized the globe [4]. Due to evolutionary selection, edible weeds thrive in places where humans disrupt the soil and they are more tolerant of environmental extremes than most commercial crops [5–7]; climate change may select for weeds and for herbicide-resistant weeds [8].

Edible weeds may be a source of interesting culinary ingredients in the global north [9] and also an important source of nutrition during food shortages [10,11]. Studies suggest that edible weeds contribute to household food supplies in cities around the globe [12–16]. Many of these plants are also recognized as having medicinal value as teas, supplements, and poultices [17,18]. While there is little data on foraging behavior in the United States, recent surveys suggest that a noticeable percentage of urban dwellers—representing a diverse group of ethnicities, cultures, and incomes—forage and prepare edible weeds [15,19–21]. For instance, a 2014–2015 survey of 105 foragers in Baltimore found that they collect 170 unique taxa of plants and fungi from the city and surrounding areas, with low and high income foragers collecting a greater variety of taxa and volume of plants than middle income foragers [21]. Survey respondents cited recreation, economic and health benefits, and connection to nature as their three main motivations for foraging.

Despite the growing recognition that foraged foods are a component of urban food systems and urban ecosystems, surprisingly little is known about their safety, nutritional value, or availability. Extant studies have drawn different conclusions about the safety of food—whether cultivated or “volunteers”—growing in urban environments. For instance, researchers who sampled popular edible weeds growing near urban roads and in the surrounding countryside of Bari, Italy, found that many of the collected species had high concentrations of essential vitamins and minerals, and only two species, *Amaranthus retroflexus* and *Plantago lagopus* had levels of Cd and Pb higher than the legal limit—even when growing in putatively “clean” rural areas [22]. See also [23].

There are a number of studies of the nutritional content of wild and feral foods (e.g., [24–33]). Our study contributes to understanding the potential health and nutritional impacts of urban foraging, by measuring nutrients and unhealthful contaminants (heavy metals, pesticides, herbicides, and PCBs) in six of the most abundant edible weed species harvested in highly-trafficked urban areas in the East Bay region of Northern California.

## Methods

Berkeley Open Source Food (https://forage.berkeley.edu), a research project at the University of California, Berkeley, mapped wild and feral edible plants, primarily leafy greens, in three residential areas bordering busy roadways and industrial zones in Berkeley, Richmond, and Oakland, California, during 2014 and 2015. Each area comprised approximately 9 square blocks; Table 1 describes them. According to the USDA, the areas in Richmond and Oakland are more than a mile from any shop that sells fresh produce; and the area in Berkeley is more than half a mile from such a shop. All have below average income, according to the U.S. Census.

**Table 1.** Boundaries of the study sites in Berkeley, Oakland, and Richmond, CA.

| city | bounding streets |
| --- | --- |
| Berkeley | Martin Luther King Jr. Way, California St., Dwight Way, Carleton St. |
| Oakland | 14th St., 17th St., Peralta St., Wood St. |
| Richmond | Carlson Blvd., Potrero Ave., S. 47th St., S. 43rd St. |

Teams of observers used the iNaturalist smartphone app with customized database fields to record estimates of the number of 1/2c servings of a variety of edible weeds at individual street addresses. The website for iNaturalist, an open-source citizen science database of observations of plants and animals, is https://inaturalist.org. The iNaturalist project page for the study reported here is https://www.inaturalist.org/projects/berkeley-open-source-food (last visited 19 May 2018). As of 19 May 2018, Berkeley Open Source Food had accumulated 631 observations of 52 species, made by 13 people.

Observations were geo-tagged and accompanied by representative photographs to support each observation—species identification and abundance. Some representative “voucher” samples were submitted to the University and Jepson Herbaria at the University of California, Berkeley; see table 4.

The number of 1/2c servings was estimated on an approximately logarithmic scale intended to facilitate accurate, repeatable categorization: <1, 1–5, 6–10, 11–20, 21–50, 51–100, 101–500, 501–1000, and >1000. Observations were generally made by pairs of observers conferring about each street address. Observers (faculty and students at the University of California, Berkeley) estimated the number of “visible” and the number of “available” servings. “Visible” servings are those available to a person with legal access to the property. “Available” servings are those within an arm’s reach of a public right-of-way, such as a sidewalk or road. While we do not know the intrinsic accuracy of the estimated number of servings, the measurements were reproducible: estimates made by different observers were calibrated against each other on sets of ten street addresses. After new observers received approximately an hour of training, inter-rater concordance of the estimates across pairs of observers was essentially perfect.

Soil samples were taken at 28 sites in Richmond, CA, on 18 June 2014, and in West Oakland, CA, on 28 August 2014. Samples were sent to the Soil and Plant Tissue Testing Lab at the University of Massachusetts, Amherst, for metal assays. The concentrations of Zinc (Zn), Copper (Cu), Arsenic (As), Selenium (Se), Lead (Pb), Nickel (Ni), Chromium (Cr), Cadmium (Cd), and Molybdenum (Mb) were measured. Soil samples were taken “per street address,” homogenizing sub-samples taken at 3–4 locations near the road at each address tested.

Tissue samples, collected 12 May 2015, targeted locations where soil testing had shown the concentration of metals to be highest (sample sites 28 and 29, Willow St. near 16th and 17th Streets, Oakland, CA), and included samples of plants growing through asphalt. Plant tissue samples were rinsed in tap water as if to make salad, then dried at the University and Jepson Herbaria at the University of California, Berkeley and sent to Brookside Laboratories (New Bremen, Ohio) to be assayed for metals (As, Cd, Cr, Cu, Pb, Hg, Mo, Ni, Se, Zn). Voucher specimens for each tested species were submitted (references Thomas Carlson 5001, 5002a/b, and 5003–5009).

Fresh/wet plant tissue samples collected on 21 March 2016 were rinsed in tap water, then assayed by SCC Global Services (Emeryville, CA) for vitamins, minerals, polyphenols, and some organic chemical contaminants, including PCBs, glyphosate, and multi-residue pesticides (via QuEChERS). Oxalis (*Oxalis pes-caprae*) and dock (*Rumex crispus*), two species high in oxalic acid, were assayed for oxalic acid.

## Results

Soil sample locations are listed in Table 2. Soil metals test results are listed in Table 3. Locations and species for dried tissue tests are listed in Table 4. Metals tests on dried tissue are given in Table 5. Wet tissue sample species and locations are listed in Table 6. Results of nutritional tests on wet tissue are reported per serving in Table 7 and per 100g in Table 8. The tables include values for kale, generally regarded as one of the most nutritious cultivated leafy greens. (Values for kale are from the USDA Food Composition Database, httpa://ndb.nal.usda.gov retrieved 15 July 2018.)

**Table 2.**
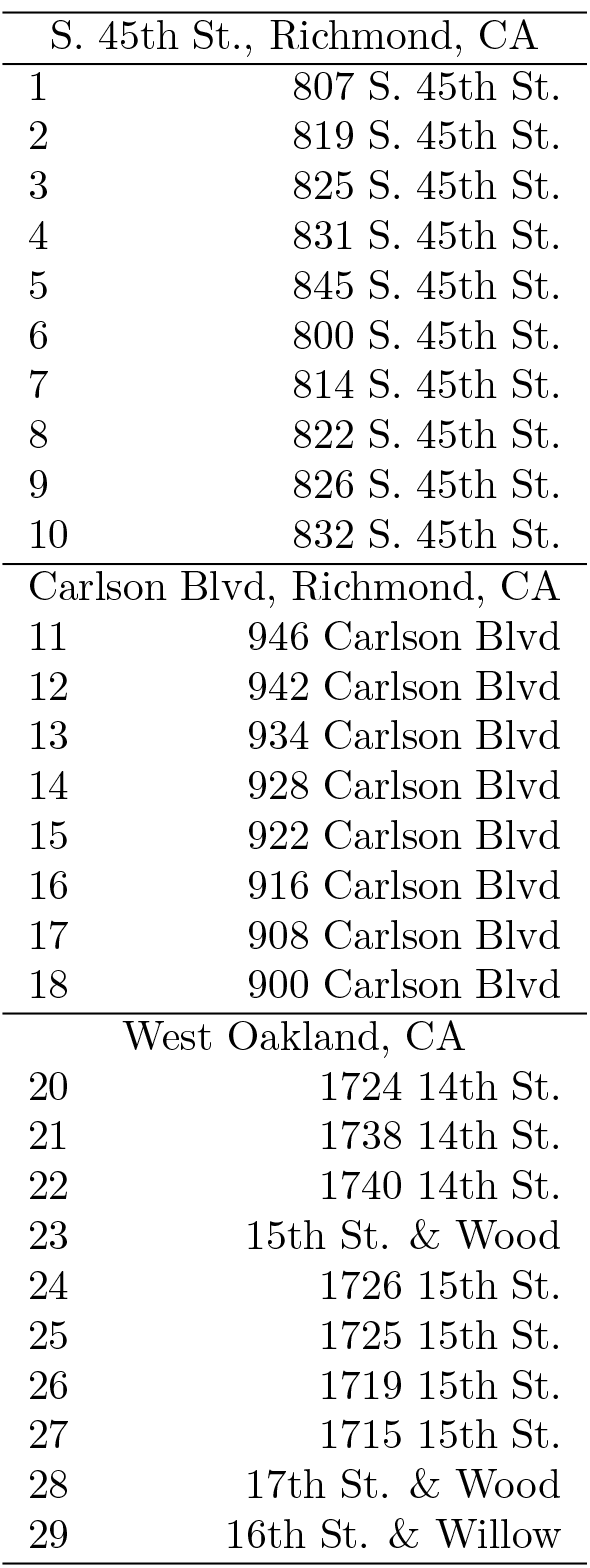
Sites where soil samples were taken. 3–4 samples were taken along the front of each street address and homogenized.

**Table 3.** Soil metal test results. See table 2 for the locations sampled. Soil tests were performed by the Soil and Plant Nutrient Testing Laboratory at the University of Massachusetts, Amherst, Center for Agriculture, Food, and the Environment

| Element | USEPA limit<br>(mg/kg) | 1-10 | 11-18 | 20-22 | 23-26 | 27 | 28 | 29 |
| --- | --- | --- | --- | --- | --- | --- | --- | --- |
| Zn | 23600 | 187.2 | 243.3 | 261.8 | 212.2 | 349.1 | 2887.2 | 453.1 |
| Cu | N/A | 41.4 | 38.6 | 40.8 | 25.6 | 37.8 | 66.8 | 63.8 |
| As | 25 | N/A | N/A | 3.4 | 1.7 | 2.8 | 5.1 | 4.1 |
| Se | 20 | N/A | N/A | 2.4 | 2.1 | 2.7 | 3.4 | 3.7 |
| Pb | 400 | 199.8 | 359.7 | 196.5 | 120.1 | 150.0 | 354.6 | 700.9 |
| Ni | 1600 | 32.4 | 30.5 | 32.7 | 23.3 | 32.3 | 73.7 | 40.9 |
| Cr | 230 | 43.7 | 35.5 | 51.3 | 39.1 | 54.9 | 56.7 | 83.6 |
| Cd | 70 | 1.2 | 0.7 | 25.7 | 21.3 | 30.9 | 58.8 | 41.9 |
| Mo | N/A | N/A | N/A | 0.8 | 0.4 | 1.0 | 0.9 | 0.7 |

**Table 4.**
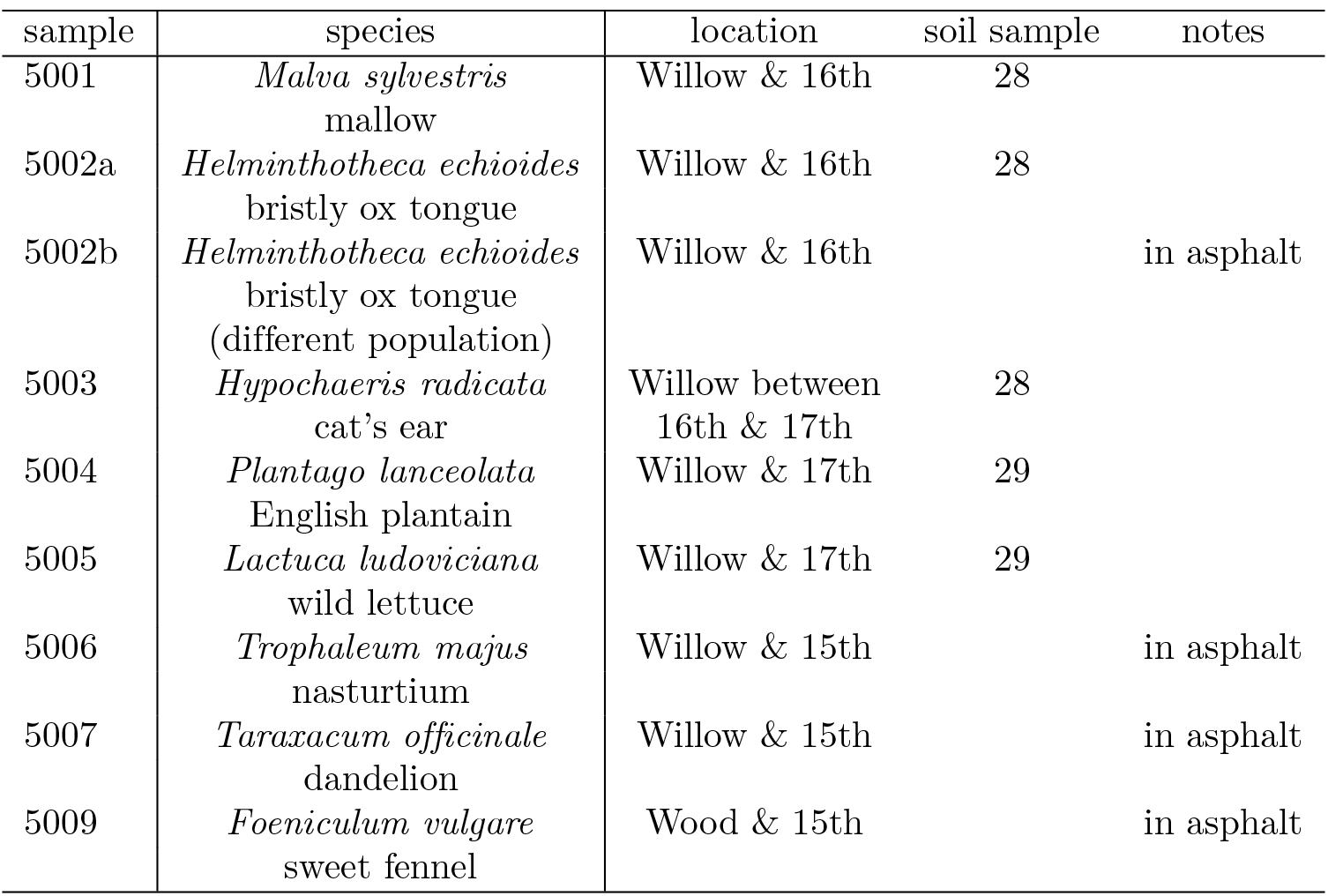
Species on which dried tissue tests were performed, and sites where samples were taken. All samples were collected on 12 May 2015 in Oakland, CA. Tissue samples were rinsed with tap water, then dried in botanical voucher sample driers at the University and Jepson Herbaria, University of California, Berkeley, before being sent for testing at Brookside Laboratories. Sample numbers refer to voucher specimens in the University and Jepson Herbaria, submitted by Prof. T. Carlson; the full reference for the first sample is, e.g., “Thomas Carlson 5001.”

**Table 5.** Metals tests of dry plant tissue. Results are in mg/kg (i.e., parts per million by mass). Tests were performed by Brookside Laboratories, Inc., New Bremen, OH. The concentration of lead (Pb) was below quantification limits in all samples. The repeated measurements differ by far more than the reported precision of the measurements, but with the exception of Cu, the measurements seem to be repeatable to two or three digits. Samples 5004-1 and 5004-2 are repeated tests on sample 5004; 5009-01 and 5009-02 were repeated tests on sample 5009. The bottom row, DWT%, gives the dry weight percentage for each sample. For instance, sample 5005 (a wild lettuce) weighed 36g before drying and 4.537g dry; the dry weight percentage is 100 × 4.537/36 = 12.6%. The original concentration of Cd in the wet tissue of 5005 is thus estimated to be 5.383 × 0.126 = 0.678ppm. See table 4 for more information about the samples.

| Element | 5001 | 5002a | 5002b | 5003 | 5004-1 | 5004-2 | 5005 | 5006 | 5007 | 5009-1 | 5009-2 |
| --- | --- | --- | --- | --- | --- | --- | --- | --- | --- | --- | --- |
| As | 1.71 | 1.69 | 1.66 | 1.68 | 1.52 | 1.61 | 1.66 | 1.62 | 1.69 | 1.59 | 1.57 |
| Cd | 0.709 | 0.859 | <0.383 | <0.391 | 0.506 | 0.500 | 5.38 | <0.383 | <0.400 | <0.392 | <0.399 |
| Cr | <0.787 | <0.767 | <0.767 | <0.781 | <0.767 | <0.769 | <0.781 | <0.766 | <0.797 | <0.784 | <0.799 |
| Cu | 13.9 | 13.6 | 7.97 | 7.87 | 8.87 | 9.04 | 17.5 | 4.79 | 8.51 | 5.27 | 5.07 |
| Pb | <3.94 | <3.83 | <3.83 | <3.91 | <3.83 | <3.85 | <3.91 | <3.83 | <3.99 | <3.92 | <3.99 |
| Hg | <0.0393 | <0.0383 | <0.0383 | <0.0390 | <0.0383 | <0.0384 | <0.0390 | <0.0382 | <0.0398 | <0.0391 | <0.0399 |
| Mo | <3.94 | <3.83 | <3.83 | <3.91 | <3.83 | <3.85 | <3.91 | <3.83 | <3.99 | <3.92 | <3.99 |
| Ni | <0.787 | <0.767 | <0.767 | <0.781 | <0.767 | <0.769 | <0.781 | <0.766 | <0.797 | <0.784 | <0.799 |
| Se | <2.36 | <2.30 | <2.30 | <2.34 | <2.30 | <2.31 | <2.34 | <2.30 | <2.39 | <2.35 | <2.40 |
| Zn | 161.0 | 69.6 | 71.8 | 110.6 | 183.1 | 183.5 | 398.2 | 115.2 | 70.9 | 30.7 | 30.5 |
| DWT% | 24.3 | 13.9 | 12.8 | 12.6 | 19.6 | 19.6 | 12.6 | 15.1 | 12.7 | 19.3 | 19.3 |

**Table 6.** Latin names and common names of species on which wet tissue tests were performed, and locations where samples were collected. Samples were collected in West Oakland, CA, on 21 March 2016. Samples were rinsed in tapwater before testing. Tests were performed by SCS Global Services in Emeryville, CA. Samples were assayed using the QuEChERS multi-pesticide residue test, which covers approximately 330 pesticides and herbicides; for glyphosate specifically; and for PCBs. No pesticides, glyphosate, or PCBs were detected in any of the samples. Nutritional test results for the samples is given in tables 7 and 8.

| species | sample location |
| --- | --- |
| chickweed | Willow & 14th St. |
| *Stellaria media* | Campbell & 14th–15th St. |
| dandelion | Willow & 14th St. |
| *Taraxacum officinale* |  |
| dock | Willow & 14th St. |
| *Rumex crispus* |  |
| mallow | Willow & Wood, 14th–17th St. |
| *Malva sylvestris* |  |
| nasturtium | Campbell & 14th–15th St. |
| *Tropaeolum majus* |  |
| oxalis, sourgrass | Willow & 15th–17th St. |
| *Oxalis pes caprae* |  |

**Table 7.**
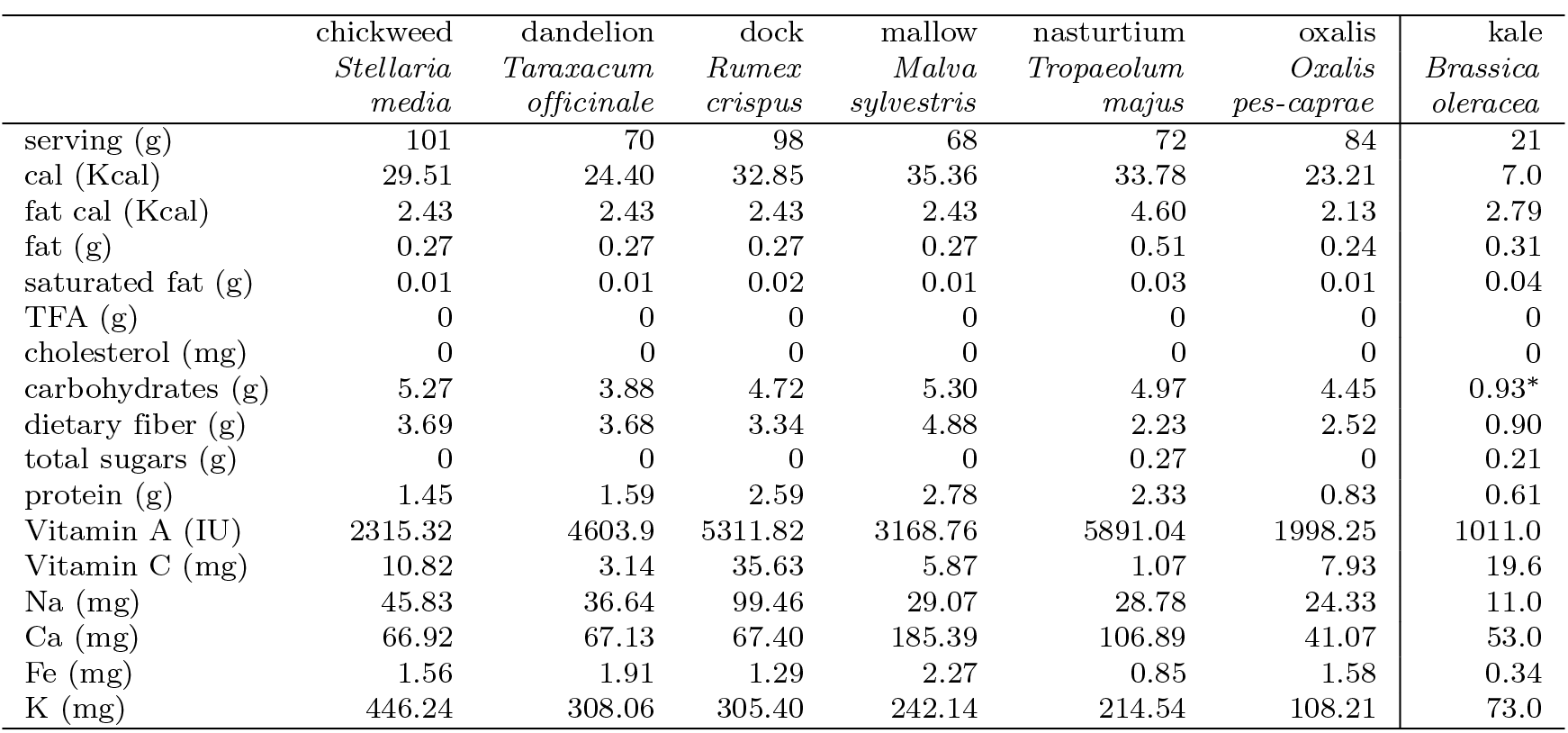
Nutritional tests of wet plant tissue (performed by SCS Global Services in Emeryville, CA) collected by Berkeley Open Source Food in West Oakland, CA, and USDA National Nutrient Database values for raw kale. Serving sizes for chickweed, dandelion, dock, and kale were 1c; serving sizes for mallow, nasturtium, and oxalis were 1/2c. Masses are listed. “cal” and “Kcal” stand for kilocalories (dietary calories) and “TFA” stands for trans fatty acids. See table 6 for sample sites. (*This number is suspiciously low—and values listed on other websites are generally 4–6g—but it is the value the USDA lists.)

**Table 8.** Nutritional tests of wet plant tissue (performed by SCS Global Services in Emeryville, CA) collected by Berkeley Open Source Food in West Oakland, CA, and USDA National Nutrient Database values for raw kale. Results are per 100g of wet tissue except total phenolics and oxalic acid, which are concentrations (mg/g) See table 6 for sample locations.

|  | chickweed<br>*Stellaria*<br>*media* | dandelion<br>*Taraxacum*<br>*officinale* | dock<br>*Rumex*<br>*crispus* | mallow<br>*Malva*<br>*sylvestris* | nasturtium<br>*Tropaeolum*<br>*majus* | oxalis<br>*Oxalis*<br>*pes-caprae* | kale<br>*Brassica*<br>*oleracea* |
| --- | --- | --- | --- | --- | --- | --- | --- |
| cal (Kcal) | 29.09 | 34.86 | 33.37 | 52.14 | 46.91 | 27.52 | 35.0 |
| fat cal (Kcal) | 2.40 | 3.47 | 2.47 | 3.58 | 6.39 | 2.52 | 13.41 |
| fat (g) | 0.27 | 0.39 | 0.27 | 0.40 | 0.71 | 0.28 | 1.49 |
| saturated fat (g) | 0.01 | 0.01 | 0.02 | 0.01 | 0.04 | 0.01 | 0.18 |
| TFA (g) | 0 | 0 | 0 | 0 | 0 | 0 | 0 |
| cholesterol (mg) | 0 | 0 | 0 | 0 | 0 | 0 | 0 |
| carbohydrates (g) | 5.19 | 5.55 | 4.79 | 7.81 | 6.90 | 5.27 | 4.42 |
| dietary fiber (g) | 3.64 | 5.26 | 3.39 | 7.20 | 3.10 | 2.99 | 4.10 |
| total sugars (g) | 0 | 0 | 0 | 0 | 0.37 | 0 | 0.99 |
| protein (g) | 1.43 | 2.27 | 2.63 | 4.10 | 3.23 | 0.98 | 2.92 |
| Vitamin A (IU) | 2282 | 6577 | 5396 | 4637 | 8182 | 2369 | 4812 |
| Vitamin C (mg) | 10.66 | 4.49 | 36.19 | 8.65 | 1.49 | 9.40 | 93.40 |
| Na (mg) | 45.17 | 52.34 | 101.04 | 42.87 | 39.97 | 28.85 | 53.0 |
| Ca (mg) | 65.96 | 95.90 | 68.47 | 273.39 | 148.46 | 48.69 | 254.0 |
| Fe (mg) | 1.54 | 2.73 | 1.31 | 3.35 | 1.18 | 1.87 | 1.60 |
| K (mg) | 439.82 | 440.08 | 310.24 | 357.09 | 297.97 | 128.29 | 348.0 |
| total phenolics<br>(mg/g) | 0.77 | 0.49 | 2.77 | 1.29 | 2.82 | 1.68 | NA |
| oxalic acid-soluble<br>(mg/g) |  |  | 0.18 |  | 10.94 |  |  |
| oxalic acid-total<br>(mg/g) |  |  | 0.39 |  | 15.42 |  |  |

### Metals in soils

Table 3 shows that a number of samples had levels of Pb approaching (in two locations) or exceeding (in one location) EPA limits, and levels of Cd greater than half the EPA limit in two locations. The levels of other toxic metals were generally far below EPA limits.

### Metals in plant tissue

The only preparation of the tissue samples was to rinse them in tapwater to remove soil and dust, then to air dry them.

Table 5 shows that despite the elevated levels of Pb and Cd, the measured amount of toxic metals in the samples was far below the US EPA maximum acceptable daily dose (RfD) for children and adults, with the exception of Cd in one species. The US EPA RfD for dietary Cd is 0.001mg/kg/d (the units are milligrams of cadmium per kilogram of body mass per day; see https://www.epa.gov/sites/production/files/2016-09/documents/cadmium-compounds.pdf, last accessed 3 June 2018). Thus, a 55kg (121lbs) adult would be expected to tolerate 0.055mg/d of Cd in food “likely […] without appreciable risk of deleterious noncancer effects during a lifetime.” The species with the highest concentration of Cd was *Lactuca ludoviciana*, a wild lettuce, which had 0.678ppm Cd in dry tissue (see Table 5). While it is hard to imagine someone eating 100g/d of *L. ludoviciana* (because it is intensely bitter), that would translate to 0.0678mg/d, over the daily limit for a 55kg person, and equal to the limit for a 68kg (150lbs) person.

### Pesticides and other toxins in fresh (wet) plant tissue

All wet tissue samples were rinsed it in tapwater to remove soil and dust, as if to make salad, but were not cleaned in any other way. The QuEChERS multi-pesticide residue test, which covers approximately 330 pesticides and herbicides, did not detect any pesticide residues, nor did specific tests for glyphosate and for PCBs.

### Nutrition of wet plant tissue

Tables 7 and 8 show that the wild greens are highly nutritious, with particularly high levels of Fe, Ca, K, protein, and dietary fiber.

## Conclusions

A healthy and sustainable food system requires a year-round, adequately abundant supply of nutrient-dense, safe, affordable food produced without depleting or contaminating vital natural resources: water, air and soil. While weeds are generally unwanted and unwelcome, our research suggests that they could be a helpful component of sustainable food systems since all the plants we collected had a higher concentration of most nutrients than domesticated leafy greens and—after rinsing in water—none had detectable levels of pesticides or PCBs, and their level of heavy metals per serving were below EPA reference doses, even though they were harvested from high-traffic and mixed-use areas. Our conclusions about safety are limited to the species and locations tested, but they are promising.

## Discussion

The nutrient density of the wild greens is high, and compares very favorably to commercially farmed produce. For instance, a cup of dock (*Rumex crispus*) or a half cup of Nasturtium (*Tropaeolum majus*) contains more than the adult RDA of Vitamin A (5000 lU). The amount of Ca per cup of mallow (*Malva sylvestris*) is almost 27% higher than the same volume of whole milk (370mg versus 276mg) and contains 72% of the protein (5.56g versus 7.69g). (Values for milk are from the USDA National Nutrient Database, https://ndb.nal.usda.gov/ndb/foods/show/01211, last visited 21 October 2018.) Per serving and per 100g, the wild greens are generally more nutritious than kale, although kale does stand out in its value of Vitamin C per 100g. While kale has higher concentration of Vitamin C than the species we tested, we did not test any wild greens in the same family as kale (*Brassicacaea*), such as wild mustard (*Hirschfeldia incana, Brassier, nigra, Sinapis arvensis*, et al.) or wild radish (*Raphanus raphanistrum*), which are also abundant in the study areas. We suspect they would have Vitamin C levels closer to that of kale.

Laboratory tests for substances toxic to humans focused on metals and chemicals expected in the study zone; there is a possibility that other contaminants were missed by the testing, such as pathogenic microbes. Other studies, e.g., [34], suggest that appropriate washing suffices to mitigate the risk of pathogenic microbes such as *E. coli* O157:H7. We did not measure the concentration of some naturally occurring minerals and chemicals in plants (e.g., phytates) that in high concentration could have negative health consequences for some humans. We did measure oxalic acid in two species known to have high concentrations.

We did not measure soil pH. Plant uptake of toxic metals varies by species and is influenced by pH, salinity, and other soil properties; there is typically a “plateau” effect that limits the ability of plants to accumulate metals as the concentration of metals in soil increases [35]. However, there is evidence that the phytoaccumulation of toxic metals in leafy greens is relatively unaffected by pH [36].

Measuring the year-round availability and abundance of these plants was beyond the scope of this study. However, some of the neighborhood mappings occurred in summer (August) of 2014, considered the worst drought year in California in 1200 years (see https://en.wikipedia.org/wiki/2011%E2%80%9317_California_drought#2014 Last visited 20 May 2018), a time expected to have little food, absent deliberate irrigation. Even during this low-production period, almost every street address in all three study areas had several servings of several different species, suggesting that wild edible greens are a reliable source of nutrition year-round.

According to the most recent data from the USDA Economic Research Service (2015), waste-adjusted availability of vegetables in the U.S is approximately 1.72 cups per capita per day [37], somewhat less than the recommended intake of 2-3 cups daily [38]. Waste-adjusted availability is is the sum of domestically produced vegetables and imports, less the waste that occurs throughout the food chain. Our observations suggest that wild greens can potentially contribute to nutrient security by filling in the gap between recommended and available daily servings of vegetables.

Many of these species volunteer on farms and in gardens, where such “accidental crops” may provide additional nutrition and income. A 2014 survey of 21 farms and gardens in the East Bay [39] found that of the 15 most most frequently reported “pest” plants, 11 are edible, and 9 are “culinary quality,” namely, Plantago, oxalis, mallow, bristly ox tongue, dandelion, blackberry, calendula, purslane, and hairy bittercress. Wild foods might also contribute to a healthy ecosystem by building soil organic matter, retaining water and nutrients in the soil, and reducing erosion. Wild plants may enhance biodiversity by serving as a habitat for insects and animals and other plants.

Recognizing urban foraging as a legitimate source of nutrition to promote a varied diet raises ethical, legal, and policy issues beyond the scope of this paper. [40] make a case for permitting foraging in municipal parks and public schools. Foraging is currently prohibited on most public lands in the US. For instance, the City of Berkeley Municipal Code section 12.44.020 states:

> Cutting, trimming or removal-Permit and inspection required. It is unlawful for any person to cut, trim, remove, mutilate, injure or in any way impair the growth of any tree, shrub or plant being or growing in or on any street, parking strip, public square, park or playground in the City, or to cause or permit the same to be done. … (Ord. 3380-NS §2, 1954)

The East Bay Regional Park District also prohibits taking any plant material whatsoever. Despite the fact that the parks use grazing, prescribed burning, and mechanical, chemical, and biological means to control some invasive species, including a number of edible species. (See, e.g., https://www.ebparks.org/civicax/filebank/blobdload.aspx?BlobID=23687, last accessed 16 July 2018. Listed edible species include sweet fennel, artichoke thistle, and Himalayan blackberry, among others.) park users are expressly forbidden from collecting any plants, including those the District seeks to eradicate. EBRPD Ordinance 38 [41] states:

> SECTION 804. PLANTS. No person shall damage, injure, collect or remove any plant or tree or portion thereof, whether living or dead, including but not limited to flowers, mushrooms, bushes, vines, grass, turf, cones and dead wood located on District parklands. In addition, any person who willfully or negligently cuts, destroys or mutilates vegetation shall be arrested or issued a citation pursuant to Penal Code Section 384a.

If foraging were permitted, what rules and norms would be needed to prevent over-foraging and harming existing ecosystems? To what extent can the “culture” of foragers ensure appropriate ecological stewardship? Because most of these edible plants are invasive, it is plausible that harvesting them will improve the overall ecology; however, such issues need to be addressed. Other countries where foraging is an established practice, including Scandinavian countries, have national rules and cultural norms governing foraging. (See, for instance, https://en.wikipedia.org/wiki/Freedom_to_roam, last accessed 16 July 2018.)

